# Brain2voice 2.0: High-performance voice synthesis brain-computer interface

**DOI:** 10.64898/2026.06.30.735633

**Authors:** Maitreyee Wairagkar, Aparna Srinivasan, Nicholas S. Card, Tyler Singer-Clark, Xianda Hou, Carrina Iacobacci, Lee M. Miller, Leigh R. Hochberg, David M. Brandman, Sergey D. Stavisky

## Abstract

Brain-computer interfaces (BCIs) offer a promising solution to speech loss due to neurological injury by decoding intended speech directly from brain activity. While recent BCIs have restored high-accuracy text-based communication, they fail to provide instantaneous voice output essential for the natural flow of conversation. Brain-to-voice BCIs address this gap by decoding voice directly from neural signals. However, even the state-of-the-art (SOTA) BCI-synthesized voice is not yet intelligible enough for real-world adoption. We introduce brain2voice 2.0, a new multimodal Transformer-based BCI decoder architecture capable of synthesizing highly intelligible voice from intracortical neural signals in real-time. Brain2voice 2.0 is trained on continuous and custom-tokenized acoustic targets and phoneme targets, leveraging their complementary speech information. We use self-supervised and adversarial training objectives that enhance acoustic feature quality and improve synthesis intelligibility. At each 10 ms timestep, the model causally outputs continuous and tokenized acoustic features for real-time voice synthesis as well as time-aligned phoneme predictions (raw phoneme error rate: 7%, comparable to the latest brain-to-text models). We evaluated this new approach on our prior intracortical brain-to-voice benchmark dataset (Wairagkar et al. 2025). Naive human listeners transcribed brain2voice 2.0 synthesized voice with a word error rate of 5.24%—an 8× improvement in intelligibility over previous SOTA results (43.75%). Brain2voice 2.0 demonstrates that highly intelligible real-time voice synthesis from neural signals is achievable, for the first time crossing the intelligibility threshold necessary for clinically viable brain-to-voice BCIs for people with paralysis.

## 1 Introduction

Paralysis caused by neurological injury can rob people of the ability to speak. This severely impacts their independence and agency. Restoring naturalistic speech is a top priority for people with speech loss [1]. In recent years, brain-computer interfaces (BCIs) have made significant strides towards restoring communication by decoding intended speech directly from brain activity into text (“brain-to-text”) or voice (“brain-to-voice”), essentially bypassing the neural injury [2–6].

Brain-to-text BCIs have demonstrated reliable high-accuracy speech decoding in multiple clinical trial participants [2–4, 7–9] and are being used for daily communication [10]. Other text-BCIs have decoded handwriting [11–15] and typing [16–18]. Despite restoring essential communication, text-based BCIs do not fulfill real-time conversation needs. While text-BCIs provide some real-time feedback by displaying words or characters on the screen as they are decoded, the decoded sentences are only finalized by a separate language model at the end of the utterance, at which point it can be transmitted as text or played aloud by a text-to-speech (TTS) software [10, 19]. The shortfall of this delayed communication becomes prominent when, for instance, talking to a BCI user without looking at their screen (e.g., over the phone or across the room): the conversation can go silent for long periods while the BCI user is composing their message (including speech decoding and manual post-hoc corrections), leading to interruptions and misunderstandings. Communication with immediacy is essential for continuous conversation, which involves rapid turn-taking and interrupting each other—neither of which is possible with current text-based BCIs.

Brain-to-voice BCIs can synthesize voice instantaneously to enable the natural flow of speech. Our recent SOTA brain-to-voice study [5] demonstrated that voice can be decoded directly from neural activity and synthesized by a BCI with real-time audio feedback at 25 ms latency. This enables a person with ALS not only to speak in real-time, but also to modulate the pitch and intonation of their BCI-voice—an important component of human speech. Despite this progress, the synthesized voice was not consistently intelligible: human listeners understood the BCI-voice with a word error rate of 43.75%, which is insufficient for daily communication. Direct high-fidelity intelligible voice synthesis is challenging, as it requires high-resolution neural information to capture rich and variable speech features. Due to this demand, prior BCIs using lower-resolution neural recording modalities have yielded lower intelligibility [3, 20–22]; delivering high intelligibility neurally-synthesized voice is a critical need before brain-to-voice neuroprostheses are ready for real-world adoption.

Here, we present a new speech BCI architecture, ‘brain2voice 2.0’, to synthesize intelligible voice from high-resolution intracortical neural signals^1^. This model yielded a word error rate of 5.24% with human listener transcription on our prior SOTA brain-to-voice benchmark dataset [23], substantially improving the intelligibility of BCI-synthesized voice without compromising near-instantaneous latency^2^. Brain2voice 2.0 is a multimodal Transformer-based model trained on acoustic targets—both continuous (rapidly varying) and tokenized (structured)—and phonemic targets with self-supervised learning and adversarial training objectives. This leverages complementary information from multiple target sources—all of which have human brain activity correlates—to enhance acoustic feature quality and improve voice synthesis intelligibility. The multimodal brain2voice 2.0 simultaneously outputs (1) continuous acoustic features (LPCNet features), (2) tokenized acoustic features (custom LPCNet tokens), and (3) phonemes, causally at every 10 ms timestep. The predicted phonemes have a low causal raw phoneme error rate of 7% (without a language model), comparable to recent brain-to-text models [2, 4]. Importantly, these phonemes are automatically time-aligned with exact temporal precision to attempted speech—a novel capability that could further serve as a basis for synchronous text decoding. The predicted continuous and tokenized acoustic features are both converted to audio waveform using a pretrained autoregressive LPCNet vocoder [24], resulting in intelligible real-time voice synthesis. Brain2voice 2.0 demonstrates that high intelligibility is achievable by decoding intracortical neural signals, establishing a strong foundation for future closed-loop brain-to-voice BCIs.

## 2 Related work

Brain-to-voice decoding remains relatively underexplored due to several challenges: the absence of ground-truth speech in people with speech loss, constrained signal-to-noise ratio of speech-relevant neural signals, limited training data, and the small number of chronic clinical trial participants with intracortical electrodes. To our knowledge, our prior work [5] is the only intracortical BCI study to demonstrate real-time continuous voice synthesis and has achieved the highest performance to date. As intracortical BCIs have consistently outperformed other neural recording modalities across different motor tasks [2, 4, 5, 11, 18], we focus on intracortical data to develop brain2voice 2.0.

Prior brain-to-voice studies using electrocorticography (ECoG) and stereoelectroencephalography (sEEG) are also limited, and the majority involve opportunistic brief studies with healthy speakers undergoing epilepsy monitoring, conducted offline and acausally. Only a small number of chronic BCI studies have been conducted with people with speech loss, which is our target population.

Among studies with healthy speakers, [25] was the first to synthesize continuous speech offline from ECoG. Subsequent studies [22, 26–30] synthesized single words, while [31–33] synthesized sentences, via direct decoding of acoustic features. However, the synthesized speech was largely unintelligible, with correlations between synthesized and target speech ranging from 0.45 to 0.80, and no WER was reported. [34] used a discrete neural latent space for single-word decoding, but the approach was not compatible with causal real-time synthesis. [35] proposed two parallel pipelines for acoustic features and text, fused via TTS and voice cloning, reporting a WER of 18.9% in offline acausal decoding. Despite the advantage of having ground-truth speech from multiple participants, none of these studies achieved intelligible voice synthesis suitable for practical use.

Among chronic BCI studies with clinical trial participants with speech loss, [21] synthesized a six-word vocabulary online with 80% accuracy from a person with ALS who retained overt speech ability, using 128-channel ECoG, but with a delay of 2 s after each word. [3] synthesized speech acausally at the end of each sentence from an individual with vocal paralysis using 253-channel ECoG, achieving a WER of 54.4%. [20] demonstrated closed-loop synthesis from 253-channel ECoG in a participant with anarthria, but with a synthesis latency of ∼1.7 s—exceeding the threshold for natural conversational feedback—and a WER of 58.8%. These studies fall short of the accuracy and latency requirements for practically usable voice synthesis, highlighting the need for a high-intelligibility, real-time brain-to-voice BCI system.

## 3 Methods

### 3.1 Participant and intracortical neural dataset

We use our prior dataset, the most extensive publicly available intracortical neural dataset [23], on which we demonstrated previous SOTA brain-to-voice performance [5]. This dataset was recorded from BrainGate2 clinical trial (ClinicalTrials.gov Identifier: NCT00912041) participant ‘T15’, a 45-year-old man with amyotrophic lateral sclerosis (ALS) and severe dysarthria over a 195-day period. This pilot clinical trial was approved under an Investigational Device Exemption by the US Food and Drug Administration (#G090003). Permission was also granted by the Institutional Review Boards at the University of California, Davis (#1843264 and #1950310) and Mass General Brigham (#2009P000505).

Neural activity was recorded from four microelectrode arrays (256 total electrodes) placed in the ventral precentral gyrus (Fig. 1) as T15 attempted to speak sentences cued on screen (8,489 unique trials). The dataset contains denoised threshold crossings (-4.5× RMS threshold) and spike-band power (250-5000 Hz) binned into 10 ms bins, normalized with a 10 s rolling window, causally smoothed, and log-transformed, yielding 512 neural features at 10 ms timesteps. Further signal processing and feature extraction details are described in [5]. Additionally, the dataset provides speech segmentation derived from the brain-to-voice model (onset and offset of each attempted word) and synthetic acoustic (LPCNet) features time-aligned with each neural trial.

**Figure 1:**
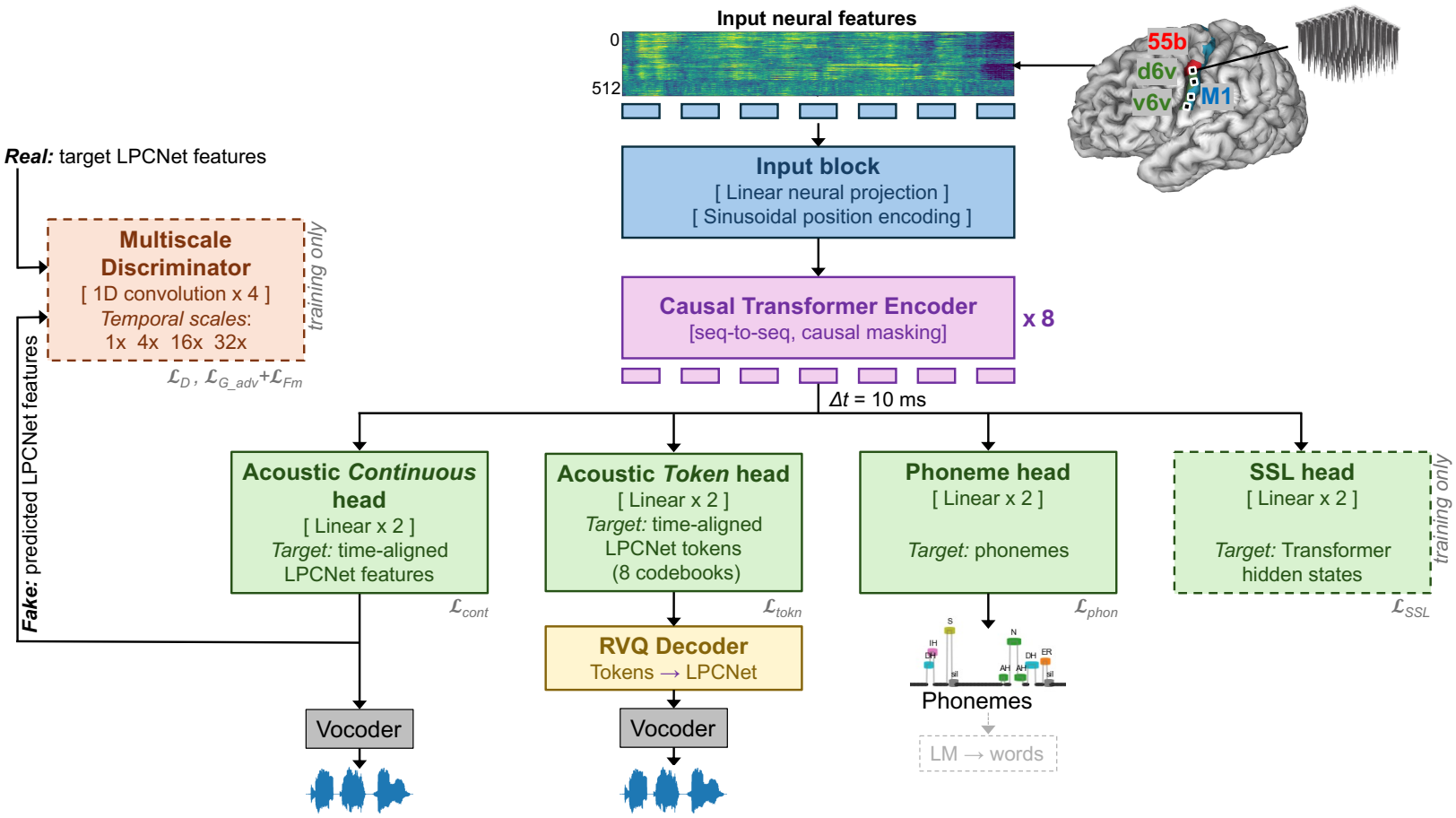
Brain2voice 2.0 neural decoder for instantaneous voice synthesis. A multimodal causal Transformer receives intracortical neural input from four microelectrode arrays in speech motor cortex of T15: ventral precentral gyrus (v6v and d6v), primary motor cortex (M1), and middle precentral gyrus (55b), of participant T15 with ALS and outputs at each 10 ms timestep: (1) continuous LPCNet features, (2) LPCNet tokens, (3) phoneme probabilities, and (4) Transformer hidden state for SSL. A multiscale discriminator provides adversarial training.

### 3.2 Brain2voice 2.0 model architecture

Brain2voice 2.0 is a multimodal Transformer-based model [36] that causally decodes continuous acoustic features, custom acoustic tokens, and phonemes at each 10 ms timestep from an input sequence of neural features (Fig. 1). The input neural features (*t*×512) are first passed through a linear projection layer followed by sinusoidal positional encoding. This projected sequence is then passed to the model backbone—a multilayer (8 layers) sequence-to-sequence Transformer encoder with causal masked self-attention (hidden size: 384, 6 heads). The resulting hidden states are passed to four parallel output heads: (1) a continuous acoustic head, (2) a tokenized acoustic head, (3) a phoneme head, and (4) a self-supervised learning (SSL) head, enabling multimodal predictions at each timestep. Additionally, a multiscale discriminator is used for adversarial training of the continuous acoustic outputs. All model hyperparameters are listed in Appendix Tables A1 and A2.

#### Continuous acoustic head

The continuous acoustic head predicts 20-dimensional continuous LPCNet features—the primary output for direct naturalistic voice synthesis. It consists of two linear layers, with rectified linear unit (ReLU) activation after the first layer. At each timestep, the predicted feature vector is converted to a 10 ms voice frame by the pretrained LPCNet vocoder [24]. Training targets are LPCNet features extracted from time-aligned synthetic speech (see Model training section).

#### Tokenized acoustic head

The tokenized acoustic head predicts LPCNet token probabilities across multiple codebooks trained using a custom tokenizer via parallel output heads, each with two linear layers with ReLU activation after the first. At each timestep, each head independently predicts a discrete token via classification. The LPCNet feature vector is reconstructed from the predicted tokens and converted to a 10 ms voice frame by the vocoder. Custom tokenizer is described below.

#### Phoneme head

The phoneme head predicts probabilities over 39 phonemes, a silence token, plus a blank token at each timestep via classification. It consists of two linear layers with ReLU activation after the first. The training target is the phoneme sequence derived from the trial text cue, providing the model with coarser-level, linguistically grounded supervision.

#### Self-supervised learning head

The SSL head predicts Transformer hidden states from masked neural inputs [37], providing auxiliary supervision beyond the acoustic and phonemic targets. Random spans of neural inputs are masked before the forward pass, and the SSL head, consisting of two linear layers with ReLU after the first, reconstructs the hidden states at masked positions, regularizing the Transformer and improving generalization

#### Multiscale discriminator

We use adversarial training with a multiscale discriminator to enhance acoustic feature quality. The discriminator operates on the continuous LPCNet features at four temporal scales: full resolution (10 ms, frame-level), 4× (40 ms, sub-phonemic), 16× (160 ms, syllabic), and 32× (320 ms, word-level), via average pooling. Each scale uses an identical four-layer 1D convolutional network with progressively expanding channels (20, 64, 128, 256), capturing local temporal patterns at increasing levels of abstraction. It produces a frame-wise realness score, with intermediate feature maps. This multiscale design promotes acoustically realistic predictions across frame, phonemic, syllabic, and word-level temporal structures critical for perceptual speech quality.

### 3.3 Custom acoustic tokenizer using residual vector quantization

This custom tokenizer quantizes continuous LPCNet features into discrete tokens, restricting the output space to a fixed tokens derived by learning representative codebooks, providing complementary information to the continuous acoustic targets. Crucially, tokenizing time-aligned LPCNet features preserves speaker-specific properties—cadence, voice characteristics, and speech patterns such as slow disarthric speech—which general-purpose tokenizers like HuBERT [38] trained on healthy speech cannot capture. These tokenizers also cannot directly decode target voice, require separate vocoder training, and do not preserve the temporal precision or latency essential for real-time closed-loop synthesis. Our custom tokenizer, by contrast, operates entirely in LPCNet feature space without latent embeddings (unlike EnCodec [39] or DAC [40]), is fully reversible and interpretable, and preserves exact temporal alignment between predicted tokens, attempted speech, and vocoder output.

The tokenizer is built using residual vector quantization (RVQ) with MiniBatch K-means clustering on LPCNet features. It consists of 8 sequential codebooks, each with 128 centroids, trained on 20-dimensional LPCNet features. Each codebook is fit sequentially to the residual after subtracting the quantized reconstruction of all previous codebooks (Fig. 2), progressively refining quantization error. The tokenizer was trained exclusively on target LPCNet features of training trials from [23], with evaluation trials held out. The tokenizer can be retrained iteratively on new data, adapting to new vocabulary or speaking style. During *encoding*, each LPCNet vector is mapped to 8 token indices—one per codebook—by sequentially assigning each residual to its nearest centroid and subtracting the corresponding quantized value. During *decoding*, the 8 token indices are mapped back to their respective centroids and summed across all codebooks to reconstruct the LPCNet features.

**Figure 2:**
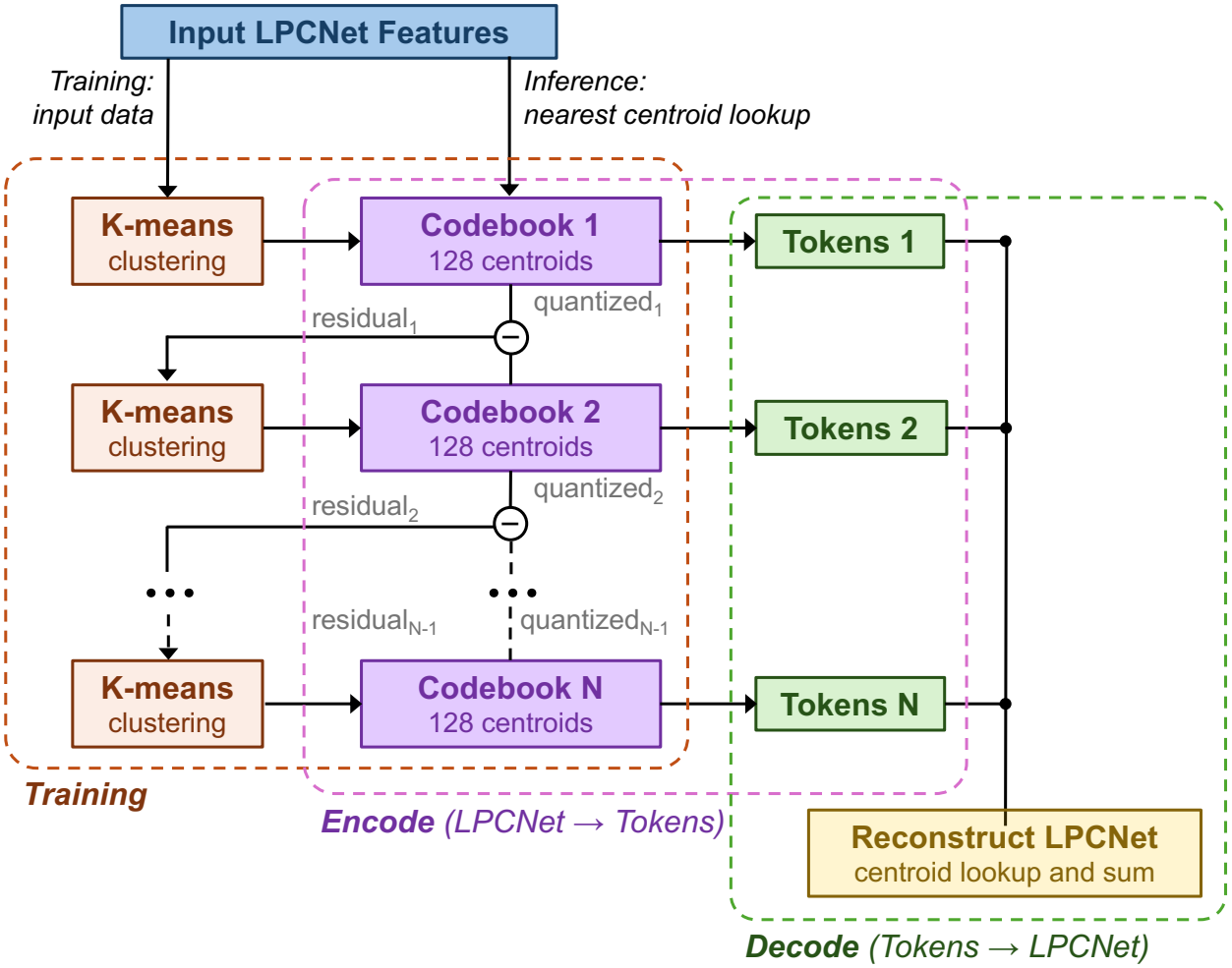
Custom LPCNet tokenizer using residual vector quantization (RVQ). The RVQ tokenizer discretizes continuous LPCNet features by learning 8 K-means codebooks sequentially by fitting residual of the previous stage. Encoding maps features to token indices via nearest-centroid lookup on successive residuals. Decoding reconstructs features by summing corresponding centroids.

### 3.4 Model training

#### 3.4.1 Target data generation

Brain2voice 2.0 is trained using three time-aligned supervised targets at 10 ms resolution: continuous LPCNet acoustic features, tokenized LPCNet acoustic features, and phoneme sequences.

##### Continuous acoustic target generation

The dataset [23] provides time-aligned synthetic LPCNet features as a proxy for T15’s intended speech. These are generated by time-stretching TTS audio of text cue to match T15’s word-level onset and offset timings—estimated from microphone recordings in the first session and from the brain-to-voice model in subsequent sessions. LPCNet features (18 spectral features, pitch period, and pitch strength) were extracted from this time-aligned audio. Here, we improved upon this pipeline by extracting LPCNet features from high quality own-voice TTS audio by cloning T15’s pre-ALS voice samples first, and time-stretching in the LPCNet feature domain via cubic interpolation, rather than stretching the raw waveform [5]. This reduces time-stretching artifacts and yields higher-quality acoustic targets.

##### Tokenized acoustic target generation

We trained the custom RVQ tokenizer on all time-aligned continuous LPCNet features from the training trials. Each continuous LPCNet feature sequence was then encoded using the trained tokenizer to obtain 8 discrete tokens (one per codebook) per 10 ms timestep, which were used as targets for the tokenized acoustic head.

##### Phoneme target generation

Phoneme sequences were obtained from trial text cues using a grapheme-to-phoneme library [41] and used directly as targets for the phoneme head. Unlike the acoustic targets, precise temporal alignment for phonemes is not required, as the phoneme head is trained using connectionist temporal classification (CTC) loss [42], handling alignment implicitly.

These three complementary targets aligned to the neural data provide the model with continuous acoustic detail, discretized acoustic structure, and high-level phonemic guidance.

#### 3.4.2 Loss functions

We use multiple primary and auxiliary losses for different output heads, that together supervise the model using complementary speech representations with distinct neural correlates:

##### Continuous acoustic loss (*L*_tokn_)

Mean squared error loss is used for continuous acoustic feature regression in unrestricted continuous output space, with upweighting for harder-to-predict high-frequency features. Since the participant’s dysarthric speech contains long silences that dilute the loss, speech timesteps are upweighted using a mask derived from target LPCNet features. The speech-to-silence weight ratio is linearly increased from 1:1 to 5:0.2 over the first 30% of training.

##### Tokenized acoustic loss (*L*_tokn_)

Cross-entropy loss is used for classifying discrete tokens for each codebook at each timestep, combined as a harmonically weighted sum—reflecting the decreasing importance of later codebooks that progressively refine smaller residuals.

##### Phonemic auxiliary loss (*L*_tokn_)

CTC loss is used for predicting phoneme logits, which does not require time-aligned phoneme targets with neural input unlike acoustic targets.

##### Adversarial training auxiliary losses

Three adversarial losses are computed on the continuous acoustic features after a warm-up period: (1) discriminator loss ( _D_) trains the discriminator to distinguish real from predicted LPCNet features using softplus loss [43] averaged across four scales; Agenerator loss (*L*_tokn_) maximizes the discriminator score on predicted outputs; (3) feature matching loss (*L*_tokn_) minimizes mean *L*1 distance between discriminator intermediate feature maps of real and predicted outputs for perceptually realistic predictions.

##### Self-supervised auxiliary loss (*L*_tokn_)

The SSL loss minimizes cosine dissimilarity between predicted and target Transformer hidden states at masked positions [37], helping the encoder learn robust representations. Targets are obtained from an unmasked forward pass.

Finally, the primary continuous and tokenized acoustic losses are combined using learned uncertainty weighting [44] because these two losses differ by an order of magnitude. Each loss is scaled by a learnable weight derived from its log variance, automatically down-weighting higher-uncertainty losses without manual tuning. The total training loss is:

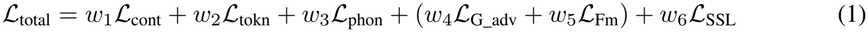

Where, *w*_1_ and *w*_2_ are tuned automatically, *w*_3−6_ are fixed (tuned manually on validation data): *w*_3_ = 0.1, *w*_4_ is linearly increased from 0.01 to 0.1 over 10,000 steps after adversarial training warm-up period, *w*_5_ = 2.0, and *w*_6_ = 0.3.

#### 3.4.3 Model training details

The model was trained using AdamW optimizer (lr = 1e-3, weight decay = 0.01, fused) with cosine decay and linear warm-up over 25,000 batches. Learnable loss weights for acoustic features were jointly optimized at lr = 1e-3 without weight decay. The discriminator used a separate AdamW optimizer (lr = 5e-5, weight decay = 0.01, non-fused) updated every alternate batch after a 5,000-step warm-up to stabilize adversarial training. Gradient clipping (max norm 1.0) and bfloat16 mixed precision were applied throughout. All components were compiled for faster training. Batches of 32 training trials each were randomly sampled and a random contiguous window equal to the context length of the model (6.4 s) was extracted from each trial. Batches were downsampled from 10 ms to 50 ms resolution (5 bins) for compute efficiency. Neural features were augmented during the last 20% of training with Gaussian noise, baseline offsets, cumulative drift noise, and 10% channel dropout to improve generalization and give an additional boost near the end. Hyperparameters were tuned manually at each development stage. 128 benchmark trials from [23] were held out for for testing and comparison with prior SOTA [5], the remaining trials were split into 30 validation trials and the rest for training. All training was performed on a single NVIDIA RTX 5090 GPU and took approximately 1 hour to complete.

#### 3.4.4 Simulated real-time inference

Brain2voice 2.0 is fully compatible with real-time closed-loop BCI decoding pipeline. We simulated real-time inference to evaluate model performance as it would operate in a closed-loop BCI. Neural input was decoded using a sliding window of 80 bins (800 ms) shifted by one 10 ms bin at each timestep—shorter than the training context (6.4 s) for faster inference. Only the last prediction from each window was retained, yielding new predictions at every 10 ms.

1. *Continuous acoustic output:* Predicted continuous LPCNet features were rescaled and passed to the pretrained vocoder to synthesize each 10 ms audio frame. This was the primary evaluation output.
2. *Tokenized acoustic output:* Predicted logits for each of the 8 codebooks were smoothed via exponential moving average, the most likely (argmax) token per codebook was selected and LPCNet features were reconstructed via the RVQ decoder, which were rescaled, causally smoothed, and passed to the vocoder at every 10 ms timestep.
3. *Phoneme output:* The most likely (argmax) phoneme was selected from predicted logits at each timestep, blank tokens were removed, and consecutive repeated phonemes were collapsed.

The model operated entirely causally—all predictions at each 10 ms timestep used only past neural data, representing the most challenging inference scenario of instantaneous voice synthesis directly translatable to real-world closed-loop BCIs.

#### 3.4.5. Evaluating intelligibility of synthesized speech

We evaluated the intelligibility of brain2voice 2.0 synthesized voice by recruiting human listeners via a crowdsourcing platform to transcribe held-out testing trials (benchmark and video sets from [23]). Each trial was independently transcribed by 7 listeners, and the median word error rate (WER) and corresponding phoneme error rate (PER) across transcriptions for each trial as in [5], were used as the primary intelligibility metrics (see A.5 for more details). We also transcribed voice samples through automatic speech recognition (ASR) via Whisper [45] to ensure consistency and scalability across conditions for comparative analyses. Additionally, we computed the Pearson correlation coefficient and mel-cepstral distortion (MCD) as objective acoustic quality metrics, following prior studies.

## 4 Results

### 4.1 Voice synthesis from intracortical neural signals

#### Quality of predicted acoustic features

High-quality voice was synthesized in real-time from continuous and tokenized LPCNet acoustic features predicted from intracortical neural data. At each 10 ms timestep, a new frame of LPCNet features was causally predicted. Fig. 3 shows the predicted continuous and tokenized LPCNet features alongside their target features, the synthesized voice waveform, and spectrograms. The predicted features closely matched the targets, with a Pearson correlation coefficient of *r* = 0.77 ± 0.06 and mean squared error (MSE) of 0.018 ± 0.005 (mean ± s.d.) across all 20 normalized features (for tokenized features, *r* = 0.68 ± 0.07 and MSE = 0.029 ± 0.007), including the harder to predict higher-frequency features that contribute to voice intelligibility.

**Figure 3:**
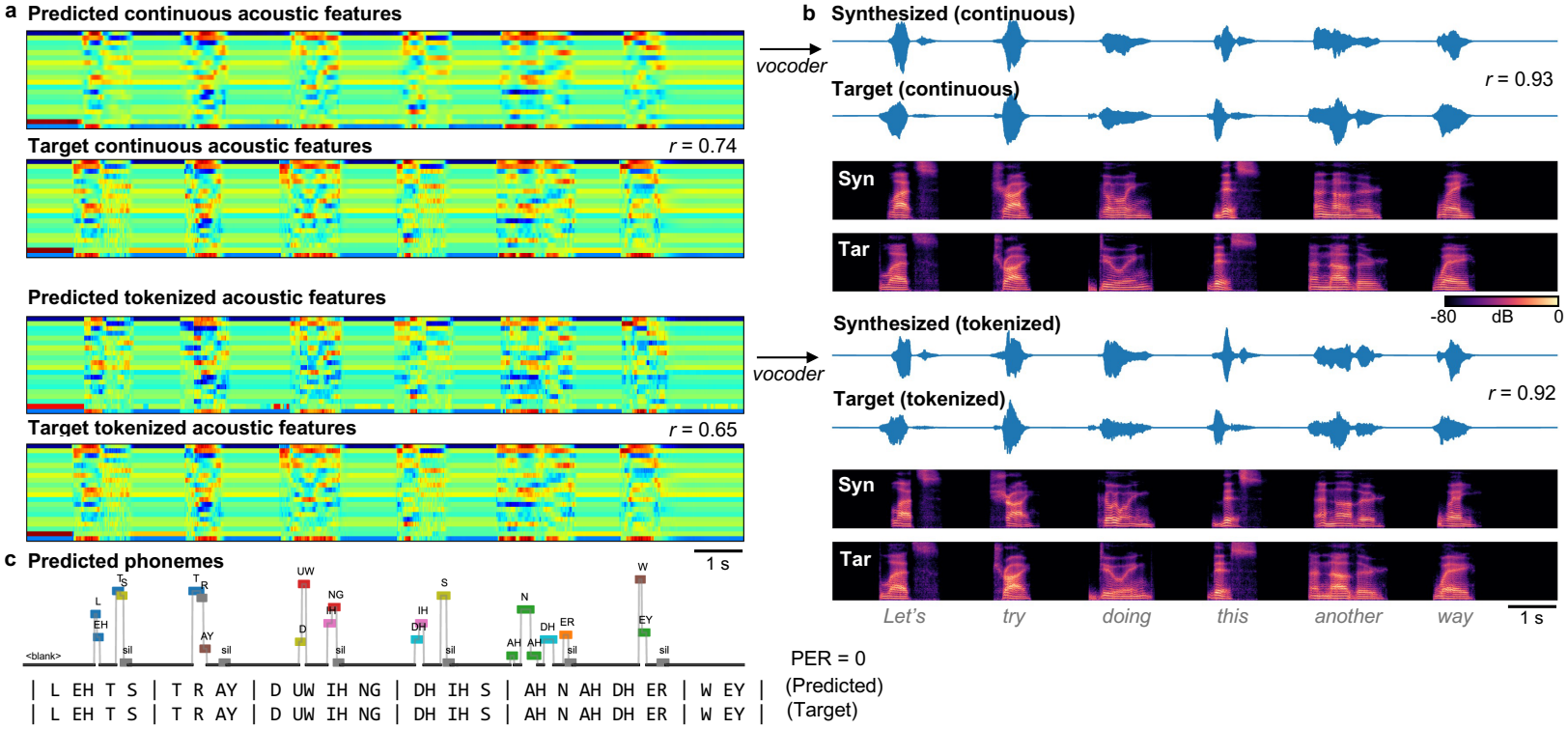
Brain2voice 2.0 acoustic and phonemic feature prediction and voice synthesis. **(a)** Predicted and target continuous (*r* = 0.74) and tokenized (*r* = 0.65) LPCNet features for an example trial. **(b)** Synthesized and target waveforms and spectrograms from continuous (r = 0.93) and tokenized (*r* = 0.92) outputs. **(c)** Predicted phoneme sequence (PER 0%) that is automatically time-aligned to the synthesized voice.

#### Intelligibility of synthesized voice

The intelligibility of brain2voice 2.0 synthesized voice as transcribed by naïve human listeners is shown in Table 1. The continuous acoustic output achieved an average WER of 5.24% and PER of 3.96%, and the tokenized acoustic output achieved WER of 5.65% and PER of 5.36% on the benchmark test trials. 79% and 74% of synthesized sentences were fully intelligible (0% WER), respectively, with a median WER and PER of 0%. Brain2voice 2.0 yielded an 8× improvement in intelligibility over the previous SOTA WER of 43.75% [5], establishing a new SOTA for brain-to-voice BCIs. Table 1 also shows intelligibility metrics on the video example set from [23] and ASR-based transcription error rates. A detailed comparison with the baseline model is provided in Appendix A.2. Example synthesized voice trials are provided alongside the target voice^3^. Appendix A.3 shows the ablation studies on the components of the brain2voice 2.0 model. Appendix A.4 shows the effect of electrode count and array location on the intelligibility of voice synthesis.

**Table 1:** Intelligibility of Brain2voice 2.0 synthesized voice.

|  | **Continuous Brain2voice 2.0** |  |  | **Tokenized Brain2voice 2.0** |  |  |
|---|---|---|---|---|---|---|
|  | Mean error rate (%) [95% CI] |  | Error-free | Mean error rate (%) [95% CI] |  | Error-free |
|  | **WER** ↓ | **PER** ↓ | trials (%) ↑ | **WER** ↓ | **PER** ↓ | trials (%) ↑ |
| **Human transcription**<br>(benchmark test set) | **5.24**<br>[2.89, 7.94] | **3.96**<br>[2.25, 5.90] | **78.89** | 5.65<br>[3.31, 8.43] | 5.36<br>[3.23, 7.67] | 74.44 |
| **Human transcription**<br>(video set) | 2.11<br>[0.26, 4.47] | 1.45<br>[0.26, 2.93] | 89.47 | 3.21<br>[0.44, 7.42] | 2.39<br>[0.49, 4.66] | 86.84 |
| **ASR transcription**<br>(benchmark test set) | 8.04<br>[5.17, 11.26] | 5.70<br>[3.66, 8.02] | 70.00 | 7.64<br>[4.84, 10.88] | 5.88<br>[3.84, 8.23] | 68.89 |
| **ASR transcription**<br>(video set) | 3.84<br>[0.26, 9.91] | 1.61<br>[0.30, 3.32] | 89.47 | 5.31<br>[1.72, 9.98] | 4.64<br>[1.66, 8.29] | 78.95 |

For comparison, the previous SOTA brain-to-voice reported in Wairagkar et al. [5], which is our baseline, achieved a WER of 43.75%, a PER of 34.00%, and 7.78% error-free trials (human transcription, benchmark test set). The full baseline comparison across evaluation sets and transcription methods is shown in Table A3.

#### Objective acoustic quality of synthesized voice

Table 2 shows objective acoustic quality metrics for continuous and tokenized synthesized voice compared to their respective target voices on benchmark trials, along with the baseline comparison. For continuous voice, Pearson correlation across 40 Mel frequencies was *r* = 0.93 ± 0.02 and MCD = 0.94 ± 0.22 dB (mean ± s.d.), with comparable results for tokenized voice (*r* = 0.92 ± 0.03, MCD = 1.10 ± 0.25 dB). These results also show substantial improvements over the prior study [5] (*r* = 0.83 ± 0.04, MCD = 2.89 ± 0.51 dB). All metrics were computed on speech periods only, with silences removed, to avoid inflating performance due to long inter-word silences in T15’s dysarthric speech. High performance across both acoustic metrics is consistent with the intelligibility results.

**Table 2:**
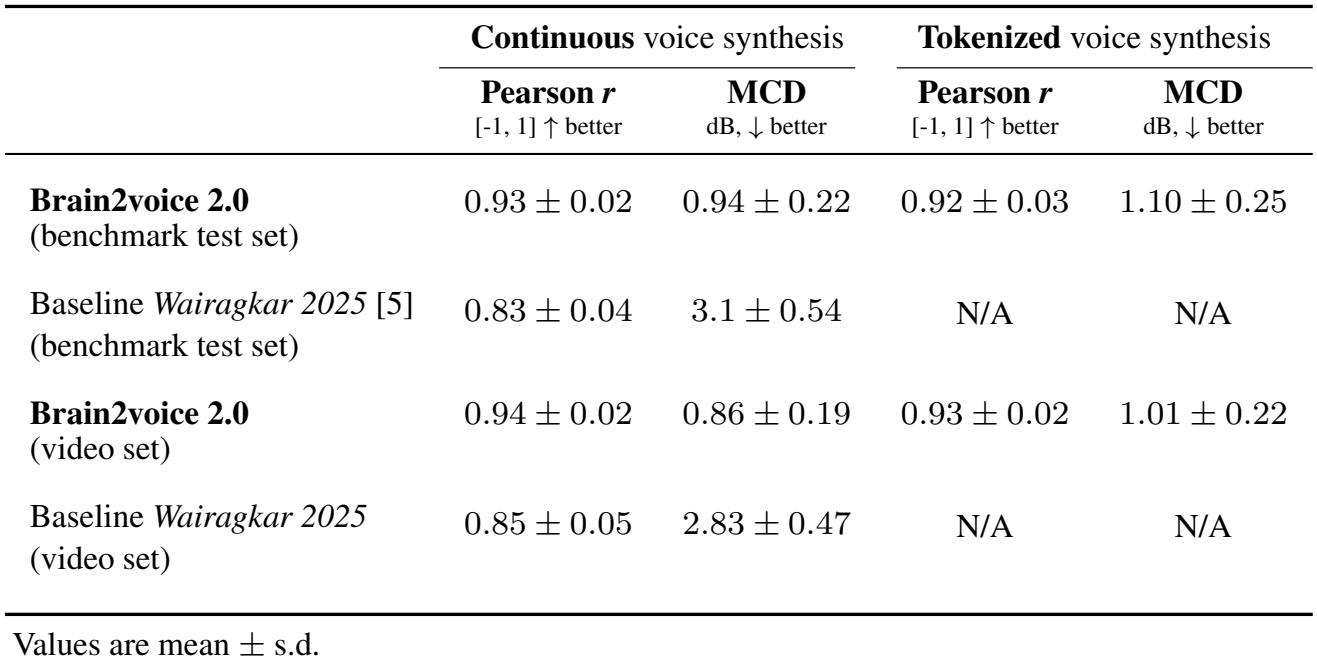
Objective acoustic quality of Brain2voice 2.0 synthesis compared to target speech.

### 4.2 Phoneme prediction from brain2voice 2.0

Brain2voice 2.0 also emitted phonemes at each 10 ms timestep, causally decoded from neural data with high accuracy. The average causal raw phoneme error rate on benchmark trials was PER = 7.03% (95% CI: [5.74, 8.39]) and on the video set was PER = 6.23% (95% CI: [3.98, 8.91]) (median PERs of 5.26% and 4.26% respectively), comparable to latest brain-to-text BCIs [2, 4]. Importantly, the predicted phonemes were automatically time-aligned with the synthesized speech and consequently with the participant’s attempted speech (Fig. 3c), without requiring explicit aligned phoneme targets. This alignment emerges automatically from the time-aligned acoustic targets used for training. This is in contrast to CTC-based brain-to-text, where phoneme alignment is only approximated by the CTC and is not precisely aligned to speech onset (this partly stems from the neural activity for each word, including preparatory activity occurring hundreds of milliseconds before speech onset).

### 4.3 Latency

Brain2voice 2.0 is suitable for real-time inference, with total computation time well under 10 ms per incoming 10 ms neural bin. Inference times were measured during simulated real-time inference on a single NVIDIA RTX 5090. Inference time for continuous acoustic LPCNet features, including feature rescaling, was 1.47 *±* 0.11 ms (mean *±* s.d.). Inference time for tokenized features was 2.11 *±* 0.21 ms, which additionally included causal logit smoothing, argmax token selection across codebooks, LPCNet feature reconstruction with the RVQ decoder, and causal LPCNet smoothing. Vocoding a single LPCNet frame took a further 1.2 ms. This leaves sufficient time for other computations required for closed-loop synthesis such as neural feature extraction, binning, normalization, smoothing, and sliding window buffering—all of which take under 4 ms as reported in [5]. Brain2voice 2.0 is therefore fully compatible with real-time closed-loop voice synthesis neuroprosthesis at 10 ms latency.

## 5 Limitations

This work used intracortical data from a single participant. While validation across multiple participants would strengthen generalizability, single-participant datasets are standard in chronic intracortical BCI research, given the significant scaling challenges involved—for example, recent high-impact chronic BCI studies all had a single clinical trial participant [2–5, 10, 11, 20, 46]. We evaluated existing data with some of the highest electrode density and signal quality available in speech-relevant brain regions. To our knowledge, no other publicly available datasets with sufficient neural signal fidelity for voice synthesis currently exist. However, it remains to be seen how performance scales with newer high-electrode count neural interfaces currently entering pilot clinical trials.

Model performance was assessed through offline evaluation with real-time inference simulation, enabling direct comparison against prior work. Offline decoding omits the potential effects of auditory feedback-driven neural adaptation by the user (which could be helpful or challenging). Evaluation in a closed-loop BCI setting remains a key next step to characterize real-world performance.

The model required ground-truth speech timing annotations from at least one seed session for initialization. Extending this to participants without precise speech onset markers, such as those with complete speech loss, is an open challenge that future work will need to address.

Finally, the dataset was collected under a structured task paradigm in which the participant attempted to speak visually-cued sentences. It remains to be seen how performance translates to self-generated spontaneous speech, as required for real-world communication.

## 6 Discussion

Brain2voice 2.0 is the most accurate and fastest voice synthesis BCI reported to date, and the first to achieve highly intelligible voice synthesis, with a human listener WER of 5.24%—an 8× improvement upon previously reported SOTA intelligibility [5]. This brings instantaneous voice communication BCIs closer to real-world viability for people with paralysis.

The model operates fully causally at each 10 ms timestep, making it suitable for future real-time closed-loop deployment. This is in contradistinction with prior voice decoding systems using ECoG and sEEG, which were neither fully causal nor real-time [3, 20, 21]. The multimodal training combining complementary targets achieved comparable intelligibility for both continuous and tokenized acoustic outputs, demonstrating that both representations successfully captured the underlying neural speech patterns. The multiscale discriminator spanning frame, phoneme, syllable, and word-level timescales improved the sharpness and perceptual quality of synthesized voice, while self-supervised learning stabilized model training by providing an auxiliary regularization.

We introduce a custom residual vector quantization-based tokenizer for LPCNet features that provides a complementary training target. It operates entirely in LPCNet feature space, is interpretable and invertible, and preserves speaker-specific voice properties and speech cadence—unlike tokenizers that operate in latent embedding space and discard speaker identity [38, 39]. This makes it particularly suitable for personalized voice synthesis. Additionally, the model’s causal time-aligned phoneme predictions could serve as a basis for simultaneous synchronous text decoding, with phoneme error rates comparable to recent brain-to-text BCIs [2, 4] despite using a fully causal architecture.

Brain2voice 2.0 arrives at a critical moment—as academic and industrial BCI programs worldwide accelerate towards clinical deployment for speech restoration. This work provides the first demonstration of high-intelligibility neural voice synthesis, offering a strong architectural foundation for next-generation voice neuroprostheses.

## Acknowledgments and disclosures

We thank participant T15 and his family and care partners for their immense contributions to this research.

## Author contributions

M.W., S.D.S., and D.M.B. conceived the study. M.W. conceptualized and designed the Brain2voice 2.0 architecture and implemented the decoder, the training algorithms, and the full framework for real-time brain-to-voice synthesis. M.W. also led the experiments, developed and coded the target speech generation algorithm, generated the cloned voice for participant T15, performed the ASR transcriptions, analyzed all data, created the figures, and wrote the manuscript. M.W. and A.S. performed the human listener evaluations. M.W., N.S.C., T.S.-C., and X.H. coded the real-time neural data collection system and built the neuroprosthetic cart system. M.W., N.S.C., and C.I. collected the primary data for this study. L.M.M. contributed to the human listener evaluations. D.M.B. led the planning and execution of the surgical implant procedure. L.R.H. is the sponsor-investigator of the multisite clinical trial. D.M.B. was responsible for all clinical trial-related activities at UC Davis. S.D.S. and D.M.B. supervised all aspects of the project. All authors reviewed and helped edit the manuscript.

## Code and data availability

Code for the Brain2voice 2.0 model and the custom RVQ LPCNet tokenizer is available at https://github.com/neuroprosthetics-lab/brain2voice2.0. The data required to train and evaluate the model are publicly available at https://doi.org/10.5061/dryad.2280gb64f.

Video and audio examples of brain2voice 2.0 synthesized speech (continuous and tokenized), drawn from the same trials that human listeners transcribed, are available alongside the target voice and ground-truth text at https://neuroprosthetics-lab.github.io/brain2voice2.0. These examples help put the reported word error rates into perspective.

## Appendix

### A.1 Brain2voice 2.0 model hyperparameters

This appendix lists the full configuration of brain2voice 2.0 implementation. Table A1 gives the model architecture hyperparameters, and Table A2 gives the training parameters, including the optimizer, learning rate schedule, loss weights, and data augmentation settings. Hyperparameters were tuned on the validation set and held fixed across all experiments.

**Table A1:**
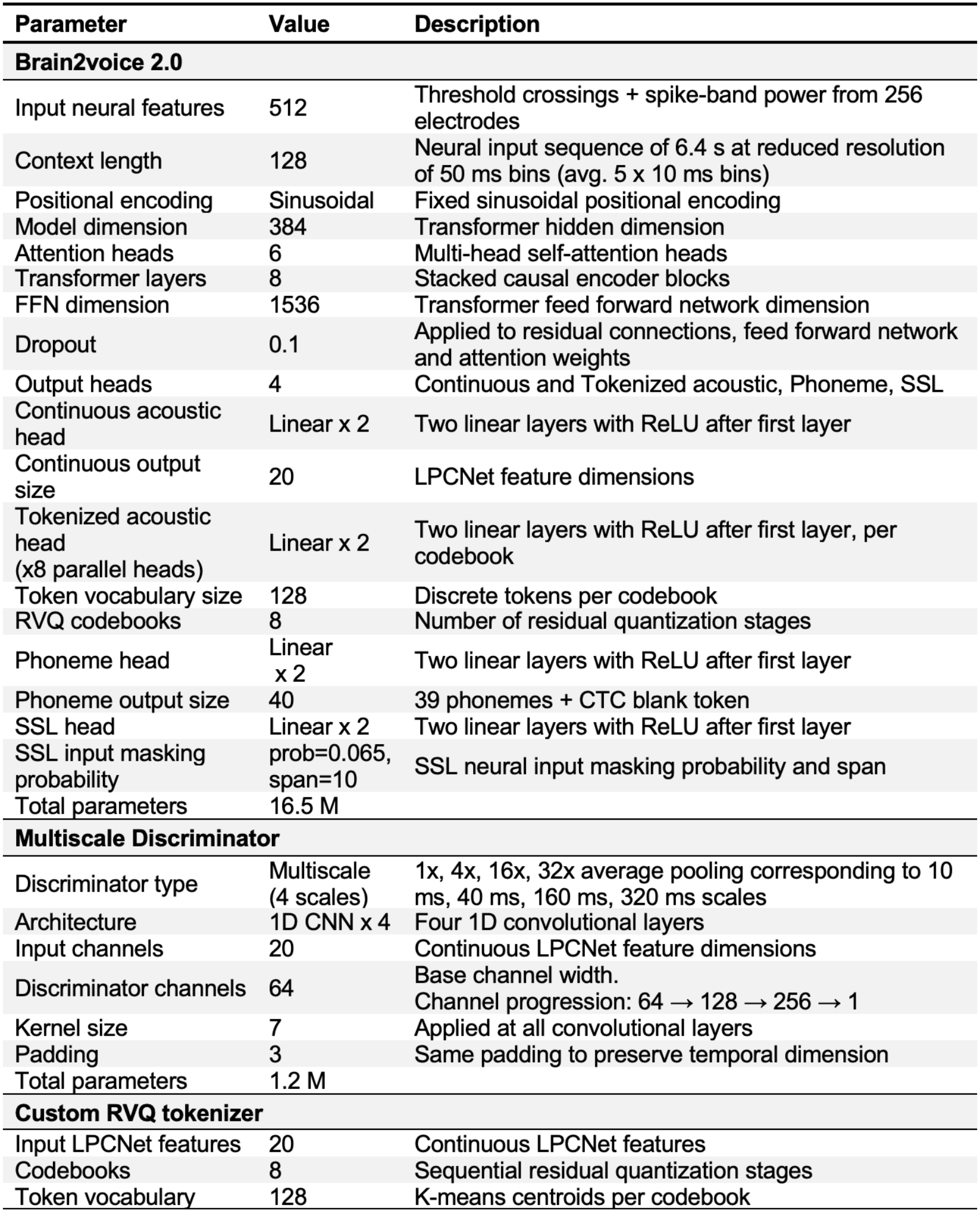
Hyperparameters for Brain2voice 2.0 model.

**Table A2:** Hyperparameters for training Brain2voice 2.0.

| Parameter | Value | Description |
|---|---|---|
| **Brain2voice 2.0** |  |  |
| Batch size | 32 | Trials per training step |
| Total training batches | 25,000 | Total training iterations |
| Max learning rate | 1×10<sup>−3</sup> | Peak learning rate after warm-up |
| Min learning rate | 1×10<sup>−5</sup> | Final learning rate at end of training |
| LR schedule | Cosine decay | Linear warm-up for first 10% of training |
| Optimizer | AdamW | Fused implementation for training efficiency |
| Weight decay | 0.01 | Applied to model parameters |
| Gradient clipping | 1.0 | Max gradient norm |
| Mixed precision | bfloat16 | Via torch.autocast |
| Data augmentation onset | 80% of training | Gaussian noise, offset, cumulative drift, channel masking |
| Learning rate adjustable loss weighting | 1×10<sup>−3</sup> | Applied to adjustable loss weighting parameters |
| Weight decay adjustable loss weighting | 0 | Applied to adjustable loss weighting parameters |
| **Discriminator training** |  |  |
| Discriminator optimizer | AdamW | Non-fused implementation |
| Learning rate | 5×10<sup>−5</sup> | Constant learning rate for discriminator |
| Weight decay | 0.01 | Applied to discriminator parameters |
| Discriminator warm-up | 5,000 steps | Discriminator is trained after 5,000 batches to allow initial Generator training stabilization |
| Gradient clipping | 1.0 | Max gradient norm |
| Discriminator update frequency | Every 2 batches | Alternating discriminator and generator updates |
| **Loss weights** |  |  |
| Continuous and token acoustic loss weight | auto | Loss weights for continuous and token heads are automatically adjusted by learned params |
| Phoneme loss weight | 0.1 | Fixed weight for phoneme CTC loss |
| Adversarial loss weight | 0.01-0.1 | Linear ramp over 10,000 steps post warm-up |
| Feature matching weight | 2.0 | Fixed weight on feature matching loss |
| SSL loss weight | 0.3 | Fixed weight on SSL loss |
| **Data augmentation** |  |  |
| Gaussian noise | σ = 0.1 | White noise added to all features at every timestep |
| Constant offset | σ = 0.05 | Baseline shift applied uniformly across trial |
| Offset scaling factor | 0.67 | Threshold crossings scaled relative to spike power |
| Cumulative drift | σ = 0.005 | Slow cumulative drift noise accumulated along time |
| Channel masking prob | 0.10 | 10% of channels are zeroed out |
| Augmentation onset | 80% | Augmentation applied only in the last 20% batches |

### A.2 Comparison with baselines

Table A3 compares brain2voice 2.0 with the baseline model from Wairagkar et al. [5] on the same evaluation sets, using both human and ASR transcription. Because the baseline model produced only continuous synthesis, tokenized results are reported for brain2voice 2.0 alone. Brain2voice 2.0 reduced WER substantially across all evaluation sets and transcription methods.

**Table A3:**
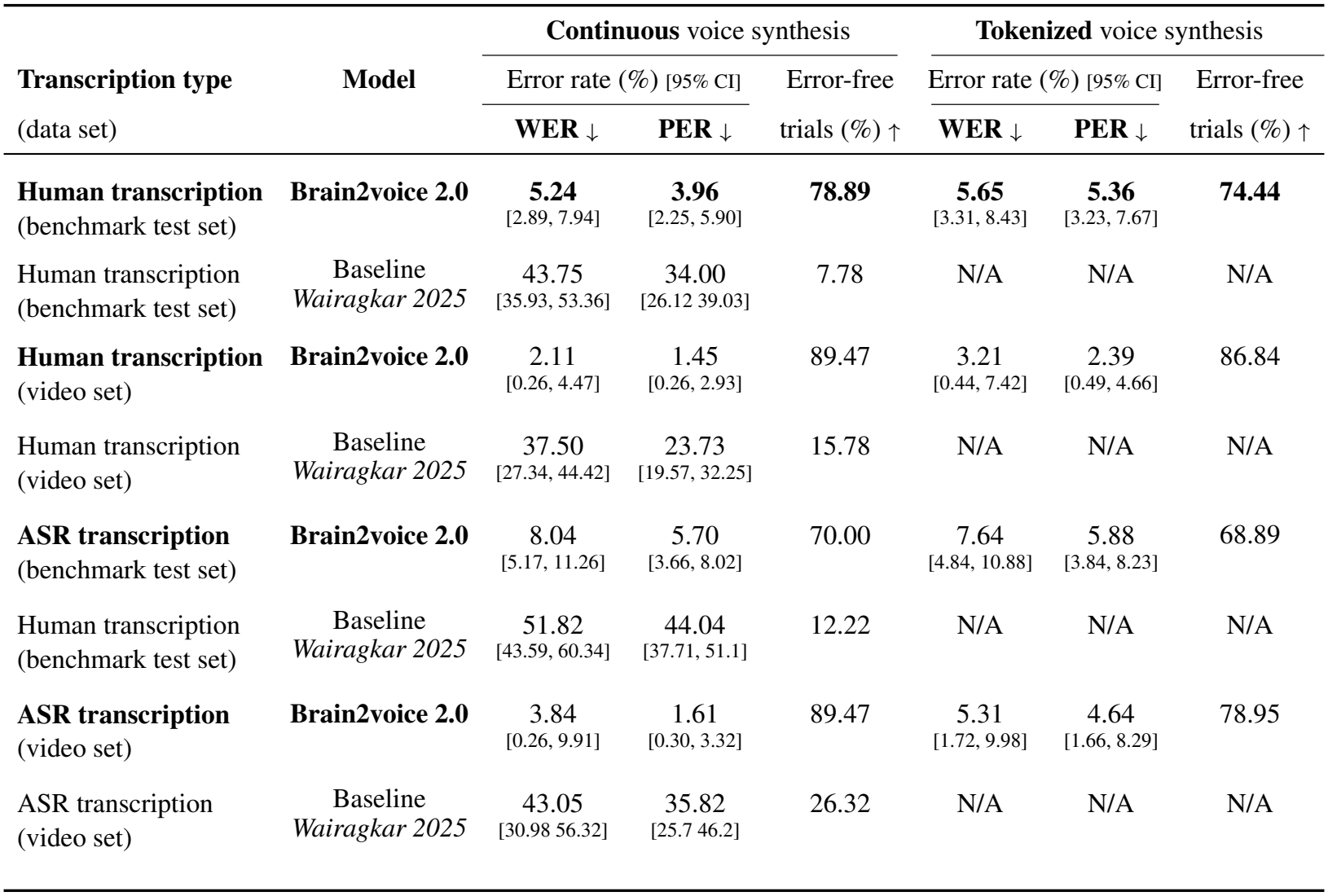
Intelligibility of Brain2voice 2.0 synthesized voice compared to baseline [5].

|  |  | **Continuous** voice synthesis |  |  | **Tokenized** voice synthesis |  |  |
|---|---|---|---|---|---|---|---|
| **Transcription type** | **Model** | Error rate (%) [95% CI] |  | Error-free | Error rate (%) [95% CI] |  | Error-free |
| (data set) |  | **WER** ↓ | **PER** ↓ | trials (%) ↑ | **WER** ↓ | **PER** ↓ | trials (%) ↑ |
| **Human transcription**<br>(benchmark test set) | **Brain2voice 2.0** | **5.24**<br>[2.89, 7.94] | **3.96**<br>[2.25, 5.90] | **78.89** | **5.65**<br>[3.31, 8.43] | **5.36**<br>[3.23, 7.67] | **74.44** |
| Human transcription<br>(benchmark test set) | Baseline<br>*Wairagkar 2025* | 43.75<br>[35.93, 53.36] | 34.00<br>[26.12 39.03] | 7.78 | N/A | N/A | N/A |
| **Human transcription**<br>(video set) | **Brain2voice 2.0** | 2.11<br>[0.26, 4.47] | 1.45<br>[0.26, 2.93] | 89.47 | 3.21<br>[0.44, 7.42] | 2.39<br>[0.49, 4.66] | 86.84 |
| Human transcription<br>(video set) | Baseline<br>*Wairagkar 2025* | 37.50<br>[27.34, 44.42] | 23.73<br>[19.57, 32.25] | 15.78 | N/A | N/A | N/A |
| **ASR transcription**<br>(benchmark test set) | **Brain2voice 2.0** | 8.04<br>[5.17, 11.26] | 5.70<br>[3.66, 8.02] | 70.00 | 7.64<br>[4.84, 10.88] | 5.88<br>[3.84, 8.23] | 68.89 |
| Human transcription<br>(benchmark test set) | Baseline<br>*Wairagkar 2025* | 51.82<br>[43.59, 60.34] | 44.04<br>[37.71, 51.1] | 12.22 | N/A | N/A | N/A |
| **ASR transcription**<br>(video set) | **Brain2voice 2.0** | 3.84<br>[0.26, 9.91] | 1.61<br>[0.30, 3.32] | 89.47 | 5.31<br>[1.72, 9.98] | 4.64<br>[1.66, 8.29] | 78.95 |
| ASR transcription<br>(video set) | Baseline<br>*Wairagkar 2025* | 43.05<br>[30.98 56.32] | 35.82<br>[25.7 46.2] | 26.32 | N/A | N/A | N/A |

#### Comparison with standard time-series architectures

Additionally, to place our results in the context of standard time-series architectures, we trained three baseline decoders (Transformer, LSTM, and CNN) to predict continuous acoustic features directly from neural data. We used continuous acoustic features because the continuous head produced the most intelligible voice output in brain2voice 2.0. Each baseline consisted of a single decoder followed by a linear output head, without the tokenized, phoneme, or SSL heads or adversarial training. The Transformer baseline uses the same encoder as brain2voice 2.0.

All models shared the brain2voice 2.0 training pipeline, including the batching protocol, context window, optimizer, learning rate, augmentation, training duration, and other training hyperparameters, and all were trained for causal, real-time synthesis. All three decoders had 8 layers and a hidden dimension of 384. The LSTM was unidirectional to preserve causality. The CNN was a causal, dilated 1D convolutional network (kernel size 3, dilation doubling per layer from 1 to 128) with left padding only, giving a receptive field of 511 timesteps. Evaluation set was also identical for all models.

Table A4 compares the performance of these common architectures. Brain2voice 2.0 outperformed all three baselines, and the Transformer outperformed the LSTM and CNN. On listening, speech synthesized by the baselines, particularly the LSTM and CNN, also had substantially lower perceptual quality.

These baseline models outperform the prior SOTA results reported in [5] because they benefit from our enhanced training pipeline. The pipeline of [5] differed substantially from that of Brain2voice 2.0, including its optimizer, loss function, context window, day-specific embedding layer, sequence-to-one training objective, and augmentation scheme. Our goal here was to test how different decoders perform within our current framework. We hypothesize that the performance of [5] would have also degraded with a LSTM or CNN decoder, as prior work has shown that Transformers outperform recurrent decoders for speech decoding [10]. Therefore [5] remains the appropriate published SOTA baseline for comparing the results of this study, as no other published model [3, 20–22] achieves comparable real-time synthesis of continuous sentences.

**Table A4:** Comparison with common baseline time-series model architectures.

| **Model type** | **Continuous** voice synthesis |  |
|---|---|---|
|  | **WER** ↓ | **PER** ↓ |
| **Brain2voice 2.0** | **8.04**<br>[5.17, 11.26] | **5.70**<br>[3.66, 8.02] |
| Transformer | 13.27<br>[9.46, 17.2] | 10.29<br>[7.26, 13.46] |
| LSTM | 18.16<br>[13.67, 22.99] | 14.33<br>[10.73, 18.23] |
| CNN | 22.42<br>[17.31, 27.91] | 18.67<br>[14.33, 23.1] |

Intelligibility was evaluated via ASR transcription

### A.3 Ablation studies on Brain2voice 2.0 model components

To understand how each component of brain2voice 2.0 contributes to intelligibility, we ran three complementary ablation studies. First, we performed a cumulative ablation, progressively adding components to a minimal model and retracing the development of brain2voice 2.0. Second, we performed a leave-one-out ablation, removing individual components from the full model to isolate each one’s contribution. Third, we performed a single-head ablation, testing a simplified model with only the continuous acoustic head. Across all ablations, we held loss coefficients, learning rate, and other hyperparameters fixed at the values tuned for the full model, since retuning each configuration separately would amount to optimizing a distinct architecture each time.

#### Ablation 1: Cumulative ablation

Table A5 presents an ablation study evaluating the contribution of each model design component of brain2voice 2.0. Starting from a baseline model with only the continuous acoustic head trained on original voice targets from our dataset in [5, 23], we progressively added components and improved voice targets. To evaluate each component’s contribution, we assessed the intelligibility of synthesized speech at each ablation via human listener transcription, where each synthesized trial (continuous and tokenized) from the benchmark set was independently transcribed by 5–7 listeners (less than 15% ablation trials were transcribed by 5 or 6 listeners, majority were transcribed by 7 listeners).

The results suggest that the auxiliary heads benefit from being trained jointly: adding the tokenized head alone increased error rates, possibly because all hyperparameters were tuned on the full model and this configuration may be suboptimal without the other components, but the subsequent addition of the phoneme head improved performance, with all three heads together yielding lower error rates. Adversarial training improved both the intelligibility and perceptual quality of continuous synthesis. Adding the SSL head helped stabilize the training and allowed the model to train longer without overfitting. Improving voice target quality by using high-quality own-voice TTS audio and reducing time-stretching artifacts through alignment in the LPCNet feature domain was equally important in improving synthesis quality. Overall, all components jointly contributed to Brain2voice 2.0, achieving the best performance through a combination of architectural choices, training improvements, and voice targets.

#### Ablation 2: Leave-one-out ablation

To isolate the contribution of each key component within the final model, we removed the tokenized head, the phoneme head, adversarial training, or the SSL head one at a time and kept everything else fixed (Table A6). For scalability, we measured intelligibility in this ablation and the next one with ASR transcription rather than human transcription. Their absolute WERs are therefore not directly comparable to Table A5.

**Table A5:** Ablation 1 - Cumulative ablation.

|  | Model configuration | Continuous<br>voice synthesis |  | Tokenized<br>voice synthesis |  | Phoneme<br>causal raw |
|---|---|---|---|---|---|---|
|  |  | **WER ↓** | **PER ↓** | **WER ↓** | **PER ↓** | **PER ↓** |
| 1 | Cont<br>*(original voice targets)* | 10.94<br>[7.5, 14.8] | 8.76<br>[6.09, 11.71] | N/A | N/A | N/A |
| 2 | Cont + Tokn<br>*(original voice targets)* | 23.17<br>[18.71, 27.97] | 21.15<br>[17.42, 24.98] | 21.50<br>[16.93, 26.39] | 18.83<br>[15.29, 22.44] | N/A |
| 3 | Cont + Tokn + Phon<br>*(original voice targets)* | 13.68<br>[10.49, 17.06] | 12.63<br>[9.59, 15.81] | 15.92<br>[12.36, 19.85] | 13.12<br>[10.35, 15.97] | 9.65<br>[8.17, 11.18] |
| 4 | Cont + Tokn + Phon<br>*(improved voice targets)* | 11.12<br>[8.14, 14.29] | 10.70<br>[7.92, 13.72] | 14.29<br>[10.9, 17.74] | 12.40<br>[9.67, 15.33] | 13.30<br>[11.58, 15.05] |
| 5 | Cont + Tokn + Phon + GAN<br>*(improved voice targets)* | 8.61<br>[5.85, 11.56] | 7.90<br>[5.56, 10.43] | 13.08<br>[9.59, 16.89] | 11.46<br>[8.39, 14.79] | 13.83<br>[12.09, 15.61] |
| 6 | Cont + Tokn + Phon + GAN + SW<br>*(improved voice targets)* | 9.28<br>[6.42, 12.33] | 8.76<br>[6.14, 11.63] | 11.81<br>[8.55, 15.38] | 10.34<br>[7.58, 13.33] | 11.73<br>[10.16, 13.39] |
| 7 | Cont + Tokn + Phon + GAN + SW + SSL<br>*(improved voice targets)* | 8.07<br>[5.06, 11.57] | 7.08<br>[4.73, 9.55] | 9.20<br>[6.22, 12.42] | 7.20<br>[4.98, 9.6] | 8.63<br>[7.11, 10.2] |
| 8 | **Brain2voice 2.0**<br>Cont + Tokn + Phon + GAN + SW + SSL<br>*(new own voice targets)* | **5.24**<br>[2.89, 7.94] | **3.96**<br>[2.25, 5.9] | **5.65**<br>[3.31, 8.43] | **5.36**<br>[3.23, 7.67] | **7.03**<br>[5.74, 8.39] |

Intelligibility of synthesized speech was assessed via human listener transcription on the benchmark test set for each ablation (mean error rate (%) [95% CI]).
*Cont*: continuous acoustic head
*Tokn*: tokenized acoustic head
*Phon*: phoneme head
*GAN*: adversarial training with multiscale discriminator
*SW*: upweighting continuous acoustic loss in speech regions
*SSL*: self-supervised learning head
*original voice targets*: time-aligned target LPCNet features from the [5, 23] dataset
*improved voice targets*: target features time-aligned by stretching in LPCNet feature domain rather than stretching target speech waveforms, to reduce time-stretching artifacts and improve acoustic quality
*new own voice targets*: target LPCNet features from high quality own-voice TTS audio cloned from T15's pre-ALS voice samples and aligned by time-stretching in the LPCNet feature domain.

Removing any single component degraded the performance and increased the WER relative to the full model. Removing SSL had the largest effect, followed by the phoneme head, the tokenized head, and adversarial training.

This result also clarifies the effect of adding the tokenized head in the cumulative ablation above, which increased the WER intermittently. However, removing this component in isolation reduces intelligibility, meaning the tokenized head helps performance. The effect of removing single components is somewhat modest. This suggests that the components in brain2voice 2.0 operate together, contributing through interactions that are difficult to isolate individually rather than through large, independent effects, complementing the findings of the cumulative ablation.

#### Ablation 3: Simplified continuous head ablation (Brain2voice 2.0 Lite)

We tested a simplified version of the model that keeps only the continuous acoustic head, removing the tokenized, phoneme, and SSL heads. We then added adversarial training, speech upweighting, and own-voice targets in sequence (Table A7; WERs were obtained from ASR transcription). Each addition improved intelligibility, and WER fell from 13.27% to 7.40%. Adversarial training, speech upweighting, and high-quality voice targets therefore each help even in a single-output configuration. This reduced variant, which we call Brain2voice 2.0 Lite, is a viable option when the added compute of a multi-head, multimodal model is undesirable, for example when simultaneous real-time phoneme decoding and quantized voice synthesis are not required.

Together, the three ablation studies show that each component is useful. Each contributes meaningfully to the best-performing configuration, and removing any single component degrades it. Because the effect of a component depends on the other components present, their contributions are not independent or additive. Rather than a single fixed architecture, brain2voice 2.0 therefore provides a modular recipe for high-accuracy, real-time, multimodal voice synthesis from intracortical signals, with the full model achieving our best human-evaluated intelligibility.

**Table A6:** Ablation 2 - Leave-one-component-out ablation.

|  | **Model configuration** | **Continuous**<br>voice synthesis |  | **Tokenized**<br>voice synthesis |  | **Phoneme**<br>causal raw |
|---|---|---|---|---|---|---|
|  |  | **WER ↓** | **PER ↓** | **WER ↓** | **PER ↓** | **PER ↓** |
| 1 | **Brain2voice 2.0** | **8.04**<br>[5.17, 11.26] | **5.70**<br>[3.66, 8.02] | **7.64**<br>[4.84, 10.88] | **5.88**<br>[3.84, 8.23] | 7.03<br>[5.74, 8.39]] |
| 2 | No Tokenized head | 11.19<br>[7.92, 14.7] | 9.08<br>[6.51, 11.89] | N/A | N/A | 11.12<br>[9.56, 14.03] |
| 3 | No Phoneme head | 11.78<br>[8.37, 15.64] | 7.85<br>[5.44, 10.43] | 11.90<br>[8.53, 15.63] | 7.36<br>[5.01 10.09] | N/A |
| 4 | No GAN | 9.86<br>[6.66, 13.39] | 5.87<br>[3.81, 8.11] | 9.51<br>[6.7, 12.6] | 5.91<br>[4.02, 7.87] | **6.85**<br>[5.54, 8.24] |
| 5 | No SSL | 12.48<br>[9.25, 15.94] | 8.14<br>[5.83, 10.65] | 13.45<br>[9.73, 17.59] | 8.58<br>[6.12, 11.23] | 10.73<br>[9.22, 12.3 ] |

Intelligibility was evaluated via ASR transcription

**Table A7:**
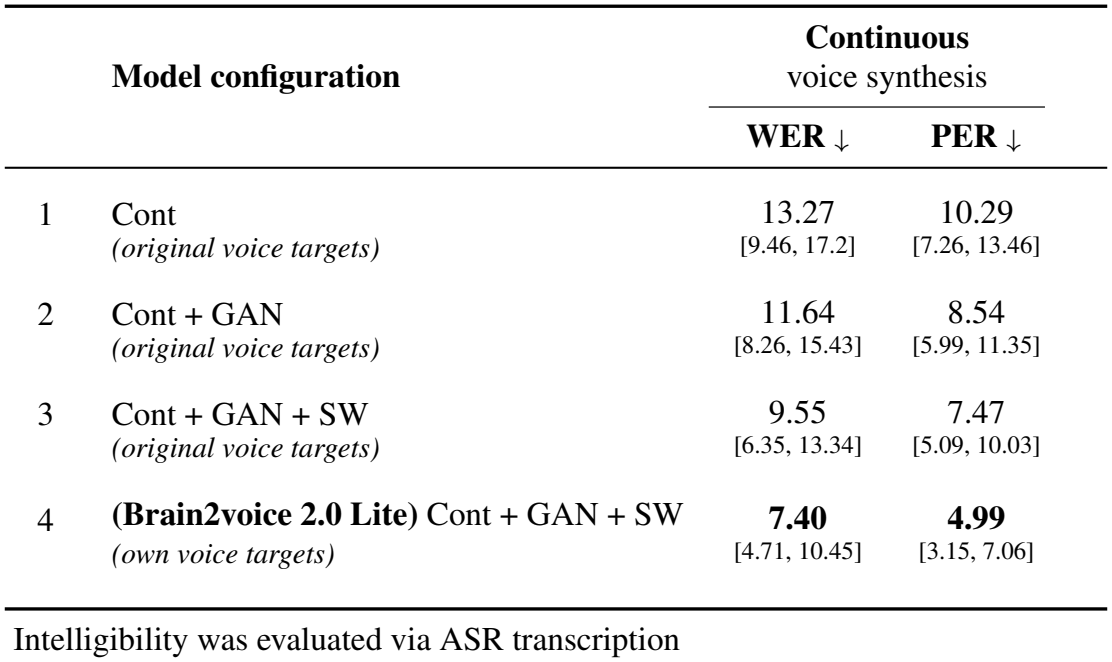
Ablation 3 - Simplified single-head ablation (Brain2voice 2.0 Lite)

### A.4 Contribution of electrode count and array location on voice synthesis intelligibility

We studied the effect of electrode count and array location on voice synthesis intelligibility. First, we randomly selected 64, 128, 192, and 256 electrodes from all arrays combined and trained separate brain2voice 2.0 models on each subset, evaluating synthesis intelligibility on benchmark trials via ASR transcription word error rate. WER decreased monotonically with increasing electrode count, though the improvement was marginal between 192 and 256 electrodes. Next, we trained separate models on each individual array to assess their relative contributions. WER was lowest for v6v, indicating it carries the highest speech-related SNR, while d6v contributed the least, yielding nearly unintelligible synthesis, consistent with prior report [4]. M1 and 55b performed intermediately, contributing more than d6v but less than v6v. These results demonstrate that voice synthesis intelligibility depends on both the number of electrodes and the speech-related SNR of the recording site, and that high number of electrodes from most relevant cortical areas are necessary for intelligible synthesis.

**Figure A1:**
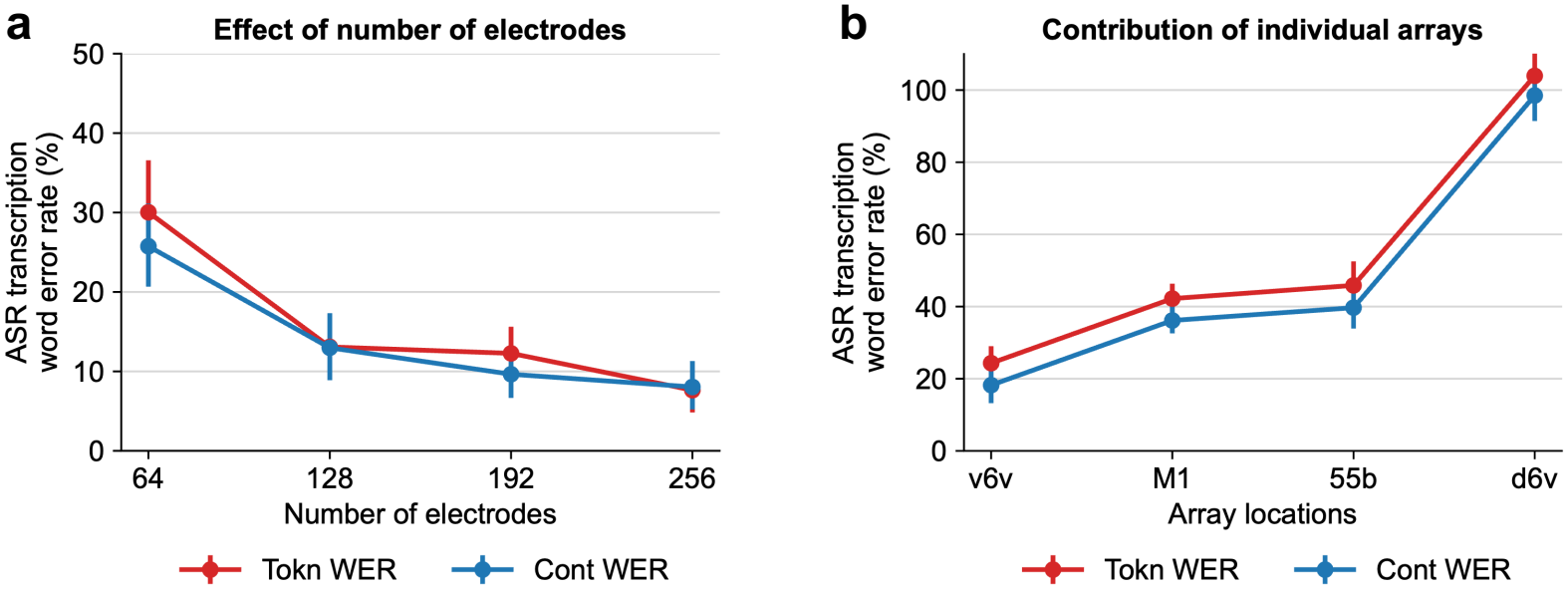
Contribution of electrode count and array location to voice synthesis. **(a)** WER on voice synthesis from models trained on randomly selected subsets of electrodes (error bars show 95% CI). WER decreases as number of electrodes increase, though the improvement becomes more marginal at higher counts. **(b)** Individual array contribution to voice synthesis measured by single-array WER. v6v contributed the most with the lowest WER, M1 and 55b contributed moderately, while d6v alone produced largely unintelligible synthesis. These results demonstrate that all four arrays together are necessary for achieving the highest possible voice synthesis quality.

### A.5 Human listener transcription of BCI-synthesized voice for evaluating intelligibility

To evaluate the intelligibility of brain2voice 2.0, human listeners transcribed each BCI-synthesized trial from the benchmark set of [5]. To allow a direct comparison with the results reported in our previous study [5], we followed the same transcription protocol: each trial was transcribed independently by 7 different listeners, and the intelligibility score for each trial was taken as the median WER of these 7 transcriptions. Taking the median across multiple transcribers is a common way to combine WERs of the same sentence [3, 5, 20].

The listeners were recruited through a crowdsourcing platform, GoTranscript^4^. Each trial was posted as a separate job with custom instructions that any transcriber from the platform’s pool, all proficient English speakers vetted by the platform, could accept and transcribe. In total, 67 unique transcribers participated, each transcribing between 1 and 132 trials (27 on average), compared to 28 transcribers who each transcribed 30–38 trials in [5]. However, both studies used an identical transcription protocol with 7 independent transcriptions per trial.

Because brain2voice [5] was originally evaluated by a different groups of transcribers, we performed two additional analyses to rule out evaluator differences as the source of improvement. First, we re-evaluated the synthesized trials from brain2voice [5] on the same crowdsourcing platform used for brain2voice 2.0. This yielded a WER of 37.5% on the same benchmark trials, consistent with the range reported in [5], indicating that transcription performance on the new platform is comparable to the previous evaluation. Second, to control for individual transcriber differences, we identified the transcribers who had transcribed both model versions. Twenty-three transcribers transcribed the same 128 sentences from both models, with brain2voice [5] always transcribed first. On these identical sentences, they had an average WER of 6.66% for brain2voice 2.0 versus 33.07% for brain2voice [5]. Together, these results show that the large improvement in intelligibility is not explained by transcriber-specific performance and reflects a genuine advance from our new approach.

#### Instructions for human listeners for transcribing BCI voice

We provided the following instructions to human listeners to transcribe the brain2voice 2.0-synthesized voice on benchmark trials:

Listen to the audio clip carefully as many times as needed and write down what you hear below each audio. Please use headphones to listen to the audio.

- Use only valid English words from the English vocabulary. Do not use any digits or numeric characters. All numbers must be written out as words (e.g., “hundred people” not “100 people”).
- The audio may be unclear, and some words may be difficult to understand. Write down the word closest to what you hear. For example, if the word is “still” and you hear something like “cell”, write “cell”.
- If you do not understand a word at all, put a ? symbol in its place.
- Ensure that the number of words you transcribe (including ? symbols) is equal to the number of words you hear in the audio.
- The transcribed sentence does not have to be grammatically or semantically correct.

## Footnotes

1 Code is available at https://github.com/neuroprosthetics-lab/brain2voice2.0.

2 Examples of video and audio of brain2voice 2.0 synthesis are available at https://neuroprosthetics-lab.github.io/brain2voice2.0.

3 Examples of video and audio of brain2voice 2.0 synthesized speech are available at https://neuroprosthetics-lab.github.io/brain2voice2.0.

4 Crowdsourcing platform used for human transcription https://gotranscript.com/.

